# CaMPARI2 Enables Stimulus-Locked Whole-Brain Activity Mapping at Cellular Resolution in Unrestrained Larval Zebrafish

**DOI:** 10.64898/2025.12.28.696792

**Authors:** Kate R. Robbins, Amelia Bredbenner, Rebecca A. Osbaldeston, Kevin S. Villafañe, Eva E. Shin, Elizaveta Merkulova, Ava Clevenger, Payton B. Delean, Cristina Campos, Graham C. Peet, Roshan A. Jain

**Author notes:** Correspondence: Roshan A. Jain.

## Abstract

Visualizing active neurons and circuits *in vivo* is critical for investigating the neural activity that underlies behavior. While several established methodologies are available to achieve this end in larval zebrafish, they are limited by the scale of tissue visualization, temporal resolution, need to restrain larvae, and/or accessibility of necessary instruments. Here, we establish a pipeline for the visualization and quantification of spatiotemporally precise whole-brain neural activity in larval zebrafish using CaMPARI2, a genetically encoded calcium indicator. Using temporally specific photoconverting UV light exposures, we capture whole-brain “snapshots” of neural activity time-locked to stimuli during unrestrained larval behavior. We optimize experimental conditions for establishing sub-second neuronal activity changes across acoustically-evoked behavioral paradigms spanning minutes to hours. We then leverage this system to pinpoint brain-wide neural activity changes during nonassociative habituation learning, observing distinct activity signatures in the subpallium, preoptic area, and habenulae that are altered through pharmacological disruption of habituation learning. This approach effectively complements the temporal precision achievable through post hoc activity detection methods and expands the accessibility of large-scale behavioral circuit dissection beyond highly specialized real-time volumetric imaging equipment.

## INTRODUCTION

Visualizing large-scale neural activity is vital for understanding how neural circuits drive and modulate behavior. In vertebrates, several established methodologies enable neuronal activity monitoring. Fluorescent activity sensors, such as genetically encoded calcium indicators (GECIs), neurotransmitter sensors, and voltage indicators, harness the transient intracellular calcium fluctuations, and membrane voltage changes associated with neural action and communication (Miyazawa et al., 2018; Sakamoto and Yokoyama, 2025). Through reversible Ca^2+^-dependent fluorescence changes, GECIs such as GCaMP and its optimized and tailored variants enable live, sensitive monitoring of activity in defined neural circuitry with high temporal resolution (Chen et al., 2013; Zhang et al., 2023). While useful for investigating cellular and subcellular calcium dynamics, reversible indicators require real-time, continuous visualization during behaviors of interest because Ca^2+^- and voltage-dependent signals are transient (Abdelfattah et al., 2019; Hiyoshi et al., 2021; Kawashima et al., 2025). High-resolution imaging in small transparent vertebrates like zebrafish can effectively capture whole brain dynamics at cellular resolution, where constraints on the visualizable tissue volume in larger organisms make it difficult to observe coordinated activity across nonadjacent neural populations (Scott et al., 2018; Vanwalleghem et al., 2018; Findling et al., 2025). These approaches typically require animal restraint or invasive implanted optical devices that may alter behavioral performance or limit how complex and generalizable recorded behaviors can be (Ahrens et al., 2012; Malvaut et al., 2020).

Post hoc detection of endogenous neuronal activity markers provides an alternative strategy that enables whole-brain activity visualization without continuous imaging. Neuronal depolarization activates the Ras-ERK signaling pathway, leading to transcription factor phosphorylation and subsequent expression of immediate early genes (IEGs) such as *Arc* and *c-fos* (Xia et al., 1996; Herdegen and Leah, 1998; Thomas and Huganir, 2004; Guzowski et al., 2005). Detection of phosphorylated ERK (pERK) or IEG expression after animals undergo a behavioral task offers a readout of recent neural activity without needing to restrain animals or limit the scale of visualizable tissue (Barykina et al., 2022; Shainer et al., 2023). Despite these advantages, activity-dependent changes captured by pERK and IEG-based approaches occur on a range of minutes to hours, limiting temporal resolution (Randlett et al., 2015; Sauvage et al., 2019; Barykina et al., 2022). Additionally, suboptimal sensitivity, weak correlation between *Arc* and *c-fos* expression and neural activity, and the differential capacity of IEGs to detect activity across neural populations complicate interpretation of which neurons were active during a given period (Fields et al., 1997; Kovács, 2008; Chiaruttini et al., 2025). Critically, these methods report a summation of all neural activity occurring within their extended detection windows, obscuring circuit activity patterns associated with distinct, temporally precise responses, behaviors, or processing stages. Although whole-brain functional calcium imaging in freely behaving animals is possible, such systems are highly specialized and broadly inaccessible due to cost and complexity (Kim et al., 2017; Scott et al., 2018; Hasani et al., 2023). Thus, accessible approaches that capture temporally specific neural activity across large tissue volumes in freely behaving animals are needed to investigate how neural circuits modulate behavior.

CaMPARI is a photoconvertable GECI that enables whole-brain capture of neural activity at cellular resolution, which can be temporally refined through precise application of photoconverting light (Fosque et al., 2015). When elevated intracellular Ca^2+^ coincides with ultraviolet (UV) light, CaMPARI undergoes irreversible green-to-red photoconversion (PC), permanently marking neurons that were active during UV exposure to yield a whole-brain “snapshot” of temporally precise activity at cellular resolution (Fosque et al., 2015; Moeyaert et al., 2018; Ebner et al., 2019). Because the ratio of red to green fluorescence scales with intracellular calcium levels, CaMPARI and its enhanced variant CaMPARI2 allow robust quantification of differential neuronal activity across large tissue volumes in diverse organisms (Fosque et al., 2015; Moeyaert et al., 2018; Edwards et al., 2020; Das et al., 2023b). Pan-neuronal CaMPARI expression in zebrafish larvae has been leveraged to detect broad total brain activity differences in fish exposed to neurotoxins and drugs (Kanyo et al., 2021, 2023; Biechele-Speziale et al., 2023). However, there is untapped potential for using brain-wide CaMPARI2 in zebrafish to characterize patterns of neural activity with finer resolution associated with temporally fixed behavioral paradigms.

Acoustically-evoked startle behavior, being both temporally stereotyped and critically modulated is a well-suited system to dissect with CaMPARI2 to understand the brain-wide neural activity patterns associated with non-associative learning in larval zebrafish. Startle behaviors like the rapid “C-start” escape responses of zebrafish are evoked by potentially threatening stimuli, and modulating when and how to respond to these stimuli are critical for survival across species (Koch, 1999; Evans et al., 2019; Zheng and Schmid, 2023). The well-characterized acoustically-evoked short-latency C-start (SLC) is driven by a pair of command-like rhombencephalic Mauthner cells that receive direct input from the statoacoustic ganglion, synapse onto contralateral spinal motor neurons, and are modulated by excitatory spiral fiber neurons as well as feed-forward and feed-back inhibitory glycinergic neurons (Burgess and Granato, 2007b; Koyama et al., 2011; Marsden and Granato, 2015; Hale et al., 2016). Repeated stimulus presentation induces robust nonassociative habituation learning, characterized by a progressive reduction in SLC initiation probability (Rankin et al., 2009; Wolman et al., 2011). This fundamental form of learning enables organisms to adapt their responses to repetitive, non-threatening stimuli, promoting behavioral flexibility (Ardiel et al., 2017; Turatto et al., 2018; McDiarmid et al., 2019). Habituation differences are a well-documented feature of diverse human neuropsychiatric conditions including autism, attention-deficit hyperactivity disorder, schizophrenia, and anxiety (Jansiewicz et al., 2004; Meincke et al., 2004; Massa and O’Desky, 2012; Sladky et al., 2012; Avery and Blackford, 2016; Jamal et al., 2021; Dwyer et al., 2023). Involvement in etiologically diverse conditions suggests a role in context-dependent behavioral modulation and learning more broadly that whole brain approaches may help elucidate (Lovett-Barron, 2021).

Here, we establish a pipeline for visualizing and quantifying spatiotemporally precise whole-brain neural activity in freely behaving larval zebrafish using CaMPARI2. Using well-defined acoustically-evoked escape behavior, we validate that CaMPARI2 captures the activation of known circuit components driving acoustically-evoked escape. We demonstrate key experimental parameters allowing CaMPARI2 to capture and highlight temporally restricted events within long experimental paradigms. We then leverage this approach to pinpoint differences in neural activity associated with different habituation states in the subpallium, preoptic region, and habenulae, which are disrupted by pharmacological blockade of NMDA-type glutamate receptors that alters the habituation state of individuals. Together, our findings position CaMPARI2 as an effective and accessible approach for resolving whole-brain, stimulus-locked, temporally specific neural activity governing behavior in unrestrained larval zebrafish.

## RESULTS

### CaMPARI2 Captures Acoustically-Evoked Activity Through UV-Dependent Photoconversion

Since CaMPARI2 photoconversion requires UV light to capture neural activity, we first assessed whether UV-dependent photoconversion could be detected when CaMPARI2 was expressed pan-neuronally in larvae. We exposed free-swimming larvae in 9 mm diameter wells to 15 3-second flashes of UV light, separated by 30 s interstimulus intervals (ISI) (**Fig. 1A**), then anesthetized and imaged whole-brain volumes with a line-scanning confocal microscope. Baseline photoconverted red signal was minimal in larvae without experimental UV exposure (**Fig 1B**), whereas larvae exposed to UV light showed green-to-red photoconversion (PC) throughout the brain, indicating widespread capture of neural activity (**Fig 1C**). To see if we could capture brain activity differences due to acoustic stimulation, we presented larvae with a similar UV exposure paradigm in which an intense 2 ms acoustic stimulus was delivered 500 ms after each UV onset, maintaining the same 22.5 s total UV exposure time across the experiment (**Fig 1D**). Larvae exposed to concurrent acoustic stimuli and UV light displayed higher levels of CaMPARI2 PC across many brain regions than larvae exposed to UV light alone (**Fig. 1C-D**), indicating that CaMPARI2 can capture acoustically-evoked neural activity in free-swimming larvae.

**Figure 1.**
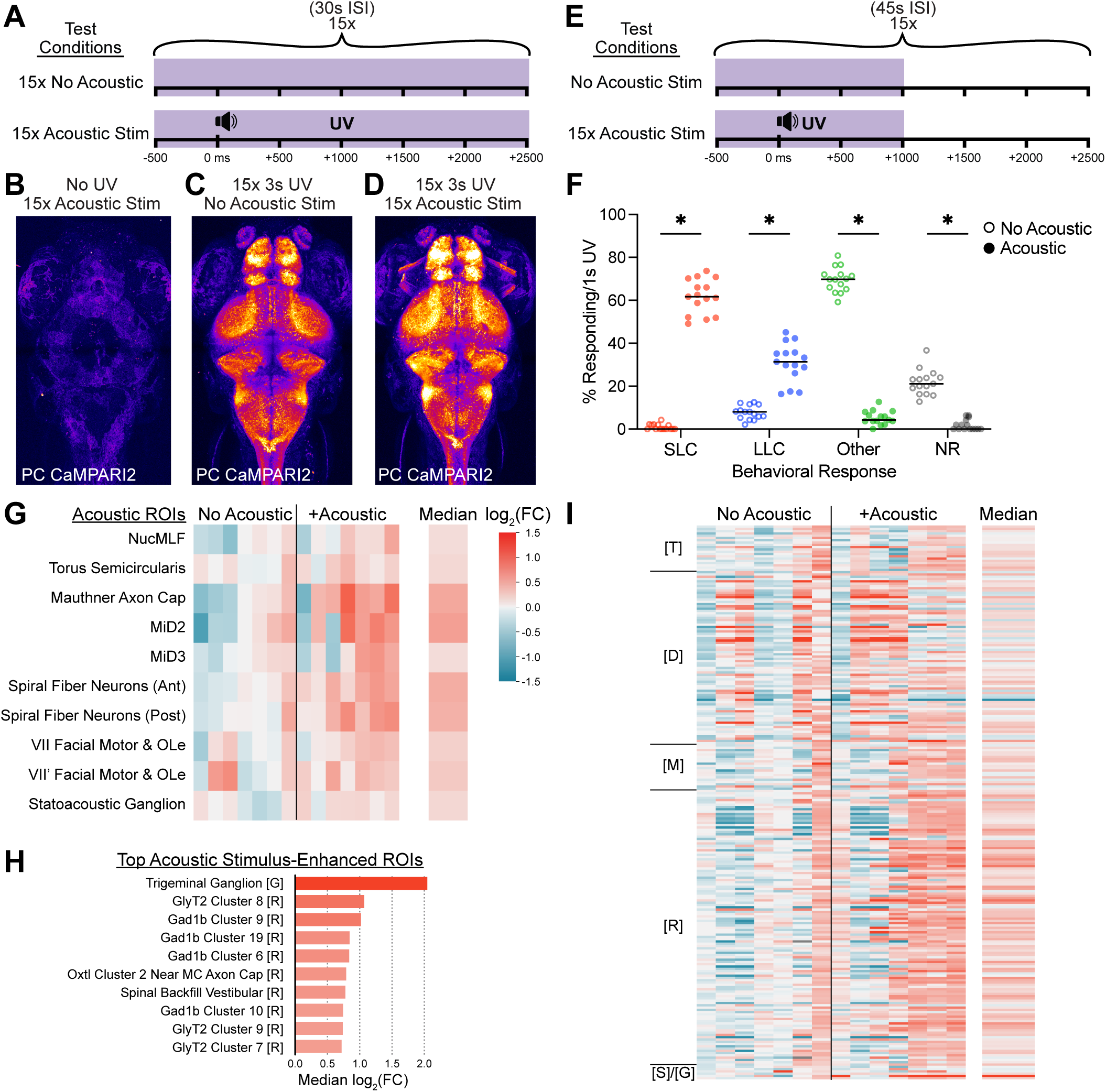
CaMPARI2 photoconversion is UV- and activity-dependent. **(A)** Schematic of UV exposure conditions for **C & D**, consisting of 15 3 s UV exposures, with or without a 2 ms acoustic stimulus 500 ms into each UV exposure. **(B-D)** Representative maximum intensity Z-projections of photoconverted CaMPARI2 from 6 dpf PTU-treated larvae expressing *HuC:CaMPARI2* that were exposed to 15 acoustic stimuli without UV exposure **(B)**, 15 UV exposure events without acoustic stimuli **(C)**, or 15 UV exposures combined with acoustic stimuli **(D)**. Photoconverted (PC) red fluorescence is presented alone, pseudocolored by intensity. **(E)** Schematic of UV exposure conditions for **F-I**, consisting of 15 1.5 s UV exposures, with or without a 2 ms acoustic stimuli presented 500 ms into each UV exposure. **(F)** Percent of tracked 6 dpf wild type TLF larvae responding to 15x 1.5s UV at 45 s ISI with no acoustic stimuli (open circles, n = 57 larvae) or with 23.5 dB acoustic stimuli (Acoustic, solid circles, n=57 larvae). Behaviors tracked over 1 s following acoustic stimuli were classified into SLC escapes (red), LLC escapes (blue), routine turns and swims (Other, green), or non-responses (NR, grey). * indicates significance by multiple Mann-Whitney tests with a 0.05 corrected p-value threshold, Holm-Šídák method. **(G-I)** Heatmaps of log_2_(fold change) in neural activity of larvae in acoustically stimulated (n = 7 larvae) and unstimulated conditions (n = 7 larvae) relative to the unstimulated condition, displaying a predetermined acoustically-active subset of ROIs **(G**), the ROIs displaying the ten largest median activity increases in the acoustically stimulated condition relative to the unstimulated condition (**H**), and the entire brain (**I**). Columns represent individual imaged under unstimulated or acoustically stimulated conditions and rows represent individual ROIs. The rightmost column shows the median log_2_(FC) across all acoustically stimulated larvae. ROIs in (**G**) were previously associated with acoustic perception, processing, or response. Bar colors in (**H**) match median log_2_(FC) legend in **(G**). ROIs in (**I**) are grouped by major brain regions along the y-axis: Telencephalon [T], Diencephalon [D], Mesencephalon [M], Rhombencephalon [R], and Spinal Cord and Ganglia [S]/[G]. ROIs with no detectable red or green fluorescence are shown in gray. Individual larvae and ROIs with >10% missing red or green fluorescence measurements were excluded from analysis.

Having validated UV- and stimulus-dependent photoconversion, we sought to increase the specificity of acoustically-evoked activity capture by scaling back the duration of each UV light exposure in our paradigm to 1.5 s (**Fig. 1E**). While relatively stereotyped and well-characterized, acoustically-evoked escape behavior can still be significantly modulated by visual input (Mu et al., 2012; Marquart et al., 2019; Martorell and Medan, 2022; Otero-Coronel et al., 2024), so we assessed larval escape behavior performance in the presence of photoconverting UV light (**Fig. 1F**). Without intense acoustic stimuli, ∼20% of larvae remained unmoving during 1 second of UV illumination (**Fig. 1F**, NR), while most larvae performed at least one movement bout during a silent 1 s UV exposure window. Larvae only very rarely initiated high-speed SLC escape maneuvers or Mauthner-independent Long-Latency C-start escape bouts (**Fig 1F**, LLC), consistent with routine turn and swim bout initiation rates observed in other illuminated contexts (Burgess and Granato, 2007a; Groneberg et al., 2020; Zúñiga Mouret et al., 2024). In contrast, introducing a single acoustic stimulus during UV illumination drastically shifted the population-level behavioral profile, with nearly all fish initiating a movement bout, most commonly an escape maneuver (**Fig 1F**, 61.7% SLC and 31.4% LLC median percentages of fish per 1 s window).

To identify which neural populations were differentially activated by acoustic stimuli and enable comparison across experimental conditions and individuals, we adapted registration and quantification pipelines originally developed for whole-brain pERK and gene expression analyses in zebrafish larvae (Randlett et al., 2015; Shoenhard and Granato, 2023). Briefly, this involved 1) registering each whole brain confocal z-stack to a previously-established reference brain in the Z-Brain Atlas (Randlett et al., 2015) 2) applying a Gaussian blur to improve regional signal consistency, 3) quantifying green and red fluorescence across the Z-Brain atlas’ 293 spatially defined neuroanatomical regions of interest (ROIs), and then 4) calculating a red-to-green ratio (RGR) for each ROI to correct for differential CaMPARI2 expression levels between ROIs and individuals. Since the neural activity captured by CaMPARI is expected to include both acoustically relevant activity, and activity associated with other simultaneous neural processes, we calculated the fold change in fluorescence in larvae exposed to both acoustic stimuli and UV light relative to larvae exposed to UV light alone for each ROI. Based on our behavioral results, we predicted that neural populations specifically involved in initiation and execution of escape behaviors would show enhanced activity in acoustically-stimulated fish. Thus we selected 10 ROIs containing neurons previously implicated in perception, processing, and/or response to acoustic stimuli (Kohashi and Oda, 2008; Hale et al., 2016; Poulsen et al., 2021; Favre-Bulle et al., 2025), and assessed their fold increase in activity relative to the matched silent photoconversion cases (**Fig. 1F**, log_2_(FC)). We excluded the ROI containing the Mauthner neurons from our analyses due to unreliable CaMPARI2 expression there in this transgenic line. While larvae exposed to acoustic stimuli displayed modest median activity increases in these acoustically-relevant ROIs, significant replicate-to-replicate variability was also evident (**Fig. 1F**). Notably, the ROI containing the trigeminal ganglion, which directly evokes SLC escape behaviors in response to tactile stimuli, showed the greatest stimulus-dependent activity increase (**Fig. 1G**) (Kimmel et al., 1990; Yokogawa et al., 2012). Other ROIs exhibiting activity increases were primarily rhombencephalic and inhibitory, consistent with extensive activation of GABAergic and glycinergic hindbrain neurons during diverse swimming behaviors (**Fig. 1G**) (Severi et al., 2018). Across the brain, an overall increase in neural activity was captured in the acoustically-stimulated condition, though we note some within-group divergence in activity patterns (**Fig. 1H**). This variability may reflect divergent behavioral profiles of the individuals in response to the 15 stimuli, or spontaneous behavioral differences in the windows prior to or following acoustically-evoked escape bouts captured by UV.

### Maximizing Temporal Restriction and Specificity in Acoustically-Evoked Activity Capture

A potential strength of the CaMPARI2 system relative to IEG-based methods is its ability to selectively capture discrete epochs of brain activity across an extended experimental time course. Because acoustically-evoked escape bouts initiate within <15 ms or ≤50 ms (for Mauthner-dependent SLC escapes or Mauthner-independent LLC escapes, respectively) and larval movement bouts typically last ≤200 ms, we restricted each UV photoconversion window to 250 ms (Burgess and Granato, 2007b; Marques et al., 2018). To maintain a similar total amount of UV exposure per fish relative to prior experiments, we exposed larvae to 100 acoustic-coupled UV exposures at 45 s ISI (**Fig. 2A**). Larvae experienced 25 s of total UV exposure distributed over a 75-minute experiment, capturing 100 behavioral epochs instead of 15. To explicitly test whether CaMPARI2 could selectively capture temporally-resolved acoustically-evoked neural activity, we assigned larvae to one of four different testing conditions, each receiving the same 100 identical acoustic stimuli varying only in the timing of 250-ms UV exposure relative to stimulus delivery (t=0 ms): -250 ms, +250 ms, +500 ms, or +750 ms (**Fig. 2A**). Because UV exposure in the -250 ms condition always concludes *before* the acoustic stimulus and subsequent escape initiation, we predicted that this condition would not capture acoustically-relevant activity. Importantly, we assessed larval behavior in response to these acoustic stimuli in the -250 ms and +750 ms experimental conditions, and found that both paradigms evoked very similar behavioral outcomes, with larvae performing mainly Mauthner-dependent SLC escapes and some Mauthner-independent LLC escapes (**Fig. 2B-C**).

**Figure 2.**
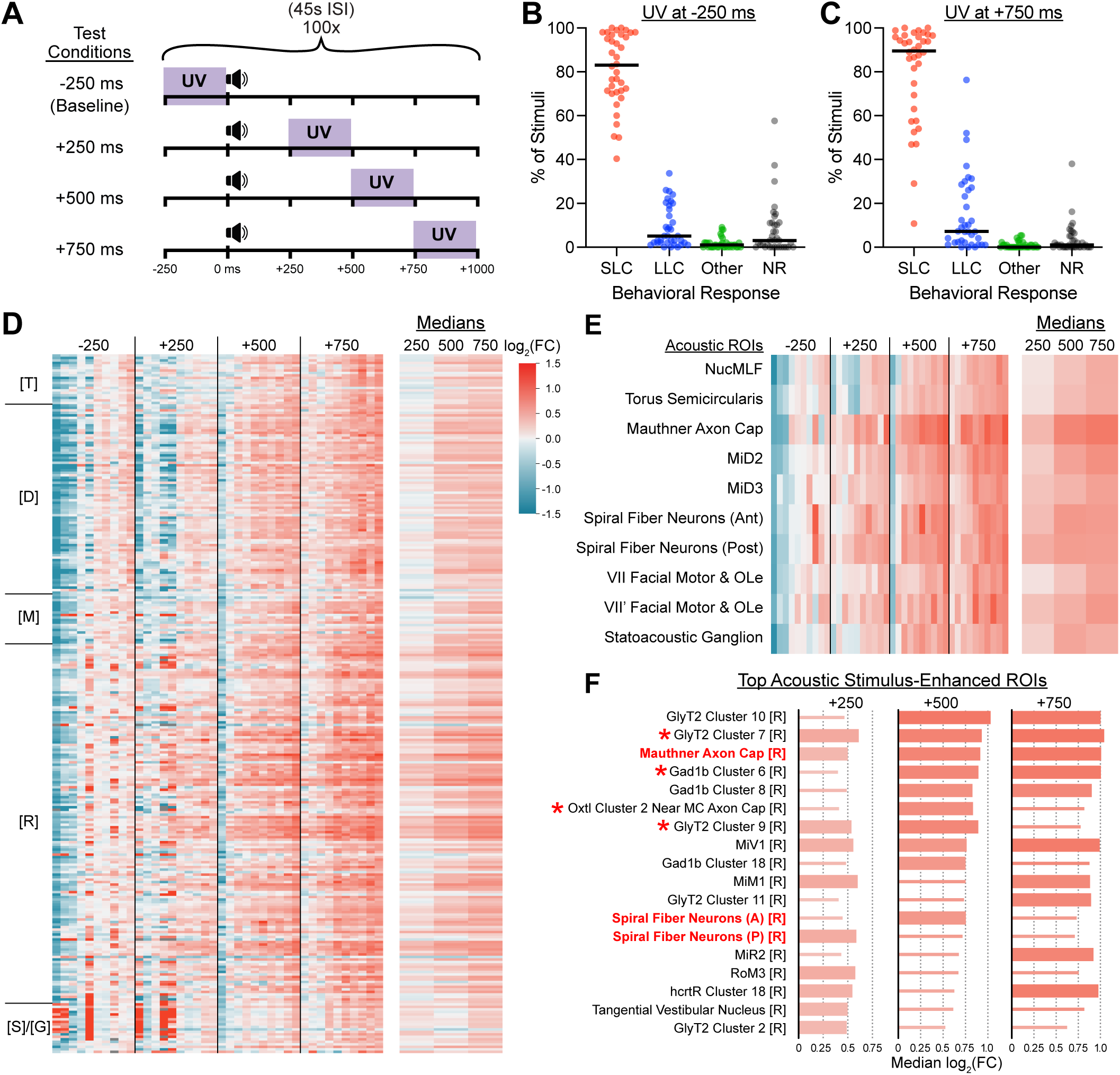
UV timing confers temporal restriction of neural activity capture by CaMPARI2. **(A)** Schematic of relative timings of acoustic stimuli and UV exposures. Each condition consisted of 100 intense acoustic stimuli presented at 45s ISI and accompanying 250-ms UV exposures beginning 250 ms before each stimulus onset (-250 ms, baseline), or 250, 500, or 750 ms after each stimulus onset (+250 ms, +500 ms, or +750 ms, respectively). **(B-C)** Frequencies of behavioral responses by 6 dpf wild type larvae to acoustic stimuli delivered according to the paradigms in (**A**) where UV was presented 250 ms prior to acoustic stimuli (**B**, n = 36 larvae) or 750ms following acoustic stimuli (**C**, n = 36 larvae). Larval behaviors were classified as Mauthner-dependent escapes (SLC, red), Mauthner-independent escapes (LLC, blue), routine turn and swim bouts (Other, green), and no response (NR, grey). **(D-E)** Heatmaps of log_2_(FC) in neural activity of larvae in pre-stimulus (-250 ms, n = 10 larvae) and post-stimulus (+250 ms, +500 ms, +750 ms, n = 10 larvae each) 250-ms UV exposure conditions relative to the pre-stimulus condition across the entire brain (**D**) and a selected subset of preselected acoustically-active ROIs (**E**). Columns represent individual larvae and rows represent individual ROIs. The rightmost columns shows the median log_2_(FC) across all larvae in each post-stimulus condition. Colors corresponding to magnitude of log_2_(FC) are indicated by color bar. grouped by major brain regions along the y-axis: Telencephalon [T], Diencephalon [D], Mesencephalon [M], Rhombencephalon [R], and Spinal Cord and Ganglia [S]/[G]. ROIs with no detectable red or green fluorescence are shown in gray. Individual larvae and ROIs with >10% missing red or green fluorescence measurements were excluded from analysis. **(F)** ROIs from (**D**) displaying the ten largest median activity increases for each post-stimulus capture group relative to the pre-stimulus condition. Thicker bars indicate the ROI was among the top 10 median log_2_(FC) value for that condition. Red text denotes ROIs known to be involved in acoustic perception, processing, or response. Asterisks indicate the ROI was among the top ten enhanced in Fig. 1H.

To identify consistent stimulus-locked activity, we calculated log_2_(FC) values relative to the median of the -250 ms condition for each ROI in each fish (**Fig. 2D**). While whole brain activity patterns of the -250 ms and +250 ms conditions displayed marked variance between replicates, within-group replicates of the +500 ms and +750 ms conditions were far more similar, with elevated activity consistently observed across individuals in many ROIs (**Fig. 2D**). Focusing on our acoustic ROI set, fold change increased in magnitude from the +250 ms to the +750 ms condition (**Fig. 2E**). We extracted the 10 ROIs with the highest fold change for each condition (**Fig. 2F**, thick bars) – all were located in the hindbrain, and several of our pre-selected acoustic ROIs now appeared among the most differentially active ROIs for at least one condition (**Fig. 2F**, red labels), alongside other descending reticulospinal clusters active during escape responses and vigorous swimming (MiV1, MiM1, MiR2, RoM3) (Gahtan et al., 2002; Lau et al., 2025). We observed substantial overlap in top-ranked ROIs between conditions such that most regions appeared among the top ROIs for multiple conditions (**Fig. 2F**), consistent with a similar behavioral profile in each condition. However, the level of activity enrichment was always highest in the +750 condition, with 50-100% higher activity captured than even the top ROI for the +250 condition (**Fig. 2F**). Taken together, introducing a 750-ms delay between acoustic stimulus delivery and UV light exposure confers sufficient temporal restriction to isolate acoustically-evoked escape behavior.

We next assessed whether further temporal restriction of the UV capture period could still efficiently detect hindbrain activity linked to escape behaviors. We thus refined our UV exposure period to 100 bouts of 125 ms UV, timing the UV relative to each stimulus as follows: -125 ms (pre-stimulus baseline), +625 ms, +750 ms, and +875 ms (all post stimulus) (**Fig. 3A**). This more restricted UV window still reliably captured widespread elevated activity in the hindbrain relative to the baseline, though broad activity in the more rostral brain areas were more muted (mesencephalon) or reduced (telencephalon) for the +750 and +875 ms conditions (**Fig. 3B**). Acoustically relevant ROIs again indicated capture of escape-related brain activity with peak median fold change values generally found in the +625 ms condition (**Fig. 3C**). Top ranked ROIs were relatively consistent between +750 & +875 ms conditions, with many consistent ROIs being detected that were also top ranked when longer UV windows were used (**Fig. 3D**, red asterisks), validating highly restricted acoustically evoked capture of acoustically-evoked escape behavior.

**Figure 3.**
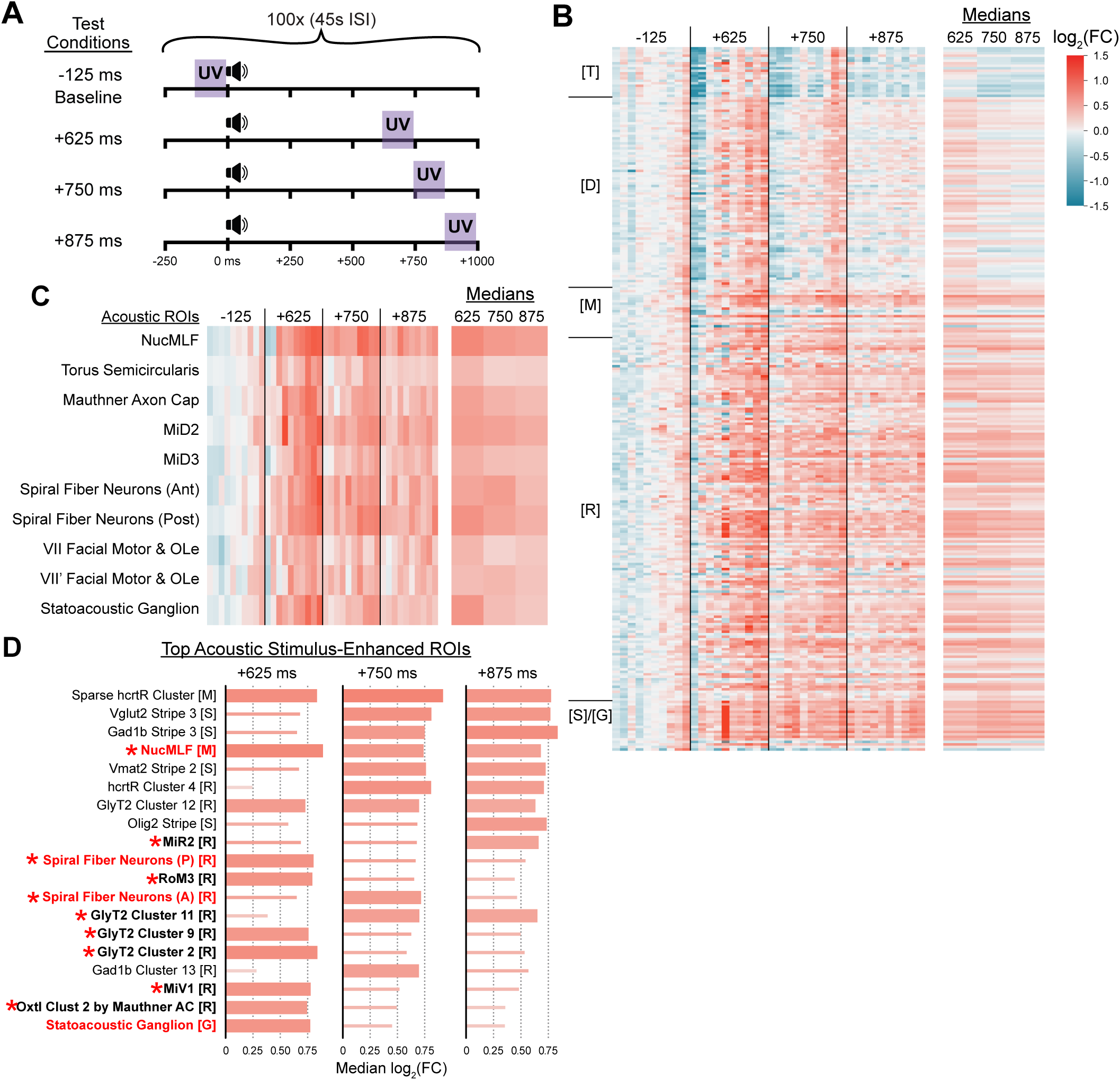
CaMPARI2 neural activity capture can be further refined through shortened photoconversion windows. **(A)** Schematic of relative timings of acoustic stimuli and UV exposures. Each condition consisted of 100 intense acoustic stimuli presented at 45s ISI and accompanying 125-ms UV exposures beginning 125 ms before each stimulus onset (-125 ms, baseline), or 625, 750, or 875 ms after each stimulus onset (+625 ms, +750 ms, or +875 ms, respectively). **(B-C)** Heatmaps of log_2_(FC) in neural activity of larvae in pre-stimulus (-125 ms, n = 10 larvae) and post-stimulus (+625 ms, +750 ms, or +875 ms, n = 10 larvae for each) 125 ms UV exposure timing conditions relative to the pre-stimulus condition across the entire brain (**B**) and a selected subset of preselected acoustically-active ROIs (**C**). Columns represent individual larvae and rows represent individual ROIs. The rightmost columns shows the median log_2_(FC) across all larvae in each post-stimulus condition. Colors corresponding to magnitude of log_2_(FC) are indicated by color bar. grouped by major brain regions along the y-axis: Telencephalon [T], Diencephalon [D], Mesencephalon [M], Rhombencephalon [R], and Spinal Cord and Ganglia [S]/[G]. ROIs with no detectable red or green fluorescence are shown in gray. Individual larvae and ROIs with >10% missing red or green fluorescence measurements were excluded from analysis. **(D)** ROIs displaying the ten largest median activity increases in post-stimulus conditions relative to the pre-stimulus condition. Bars indicate median log_2_ (fold change) value. Bolded lines indicate ROIs that appeared in the top ten for more than one post-stimulus condition. Red text denotes ROIs known to be involved in acoustic perception, processing, or response. Purple text denotes ROIs that appeared in the top ten for at least one 250-ms post-stimulus condition (Fig. 2E). **(F)** ROIs from (**B**) displaying the ten largest median activity increases for each post-stimulus capture group relative to the pre-stimulus condition. Thicker bars indicate the ROI was among the top 10 median log_2_(FC) value for that condition. Red text denotes ROIs known to be involved in acoustic perception, processing, or response. Asterisks indicate the ROI was among the top ten enhanced in Fig. 2F.

### CaMPARI Captures Differences in Habituation Learning States

Having successfully established temporal restriction in CaMPARI2 activity capture through optimized UV light delivery conditions, we leveraged the temporal control of CaMPARI2 to reveal brain-wide activity differences underlying different behavioral states of habituation. With repeated acoustic stimuli, zebrafish first shift their behavioral bias from performing Mauthner-dependent SLC escapes towards responding with Mauthner-independent LLC escapes (Jain et al., 2018). Larvae eventually discontinue motor responses to the stimuli, following conserved features of habituation learning where shorter ISIs between repeated stimuli produce faster learning and stronger behavioral attenuation in fewer stimuli than longer ISIs (Rankin et al., 2009; Wolman et al., 2011; Jain et al., 2018). We thus adapted our stimulation paradigm to capture brain activity when fish experience acoustic stimuli in a higher habituation state by pre-exposing fish to an extra 50 stimuli at 1 s ISI, which were followed by100 UV-coupled stimuli at 6 s ISI (**Fig. 4A**). This captured acoustically-evoked brain activity where larvae are more likely to perform LLCs or not respond compared to matched fish receiving only 100 UV-coupled acoustic stimuli at 45 s ISI where responses are primarily SLCs **(Fig. 4B**). Importantly, these two stimulation paradigms received identical amounts of photoconverting UV light capturing an identical number of UV-accompanied stimuli (250 ms UV exposure initiated 750 ms after each of 100 stimuli). Thus the “Low Habituating” 45s ISI condition captures the transition of naïve fish beginning to shift their bias away from SLCs, and the “High Habituating” 6s ISI condition captures brain activity after fish have already shifted their behavioral profile (**Fig. 4C**).

**Figure 4.**
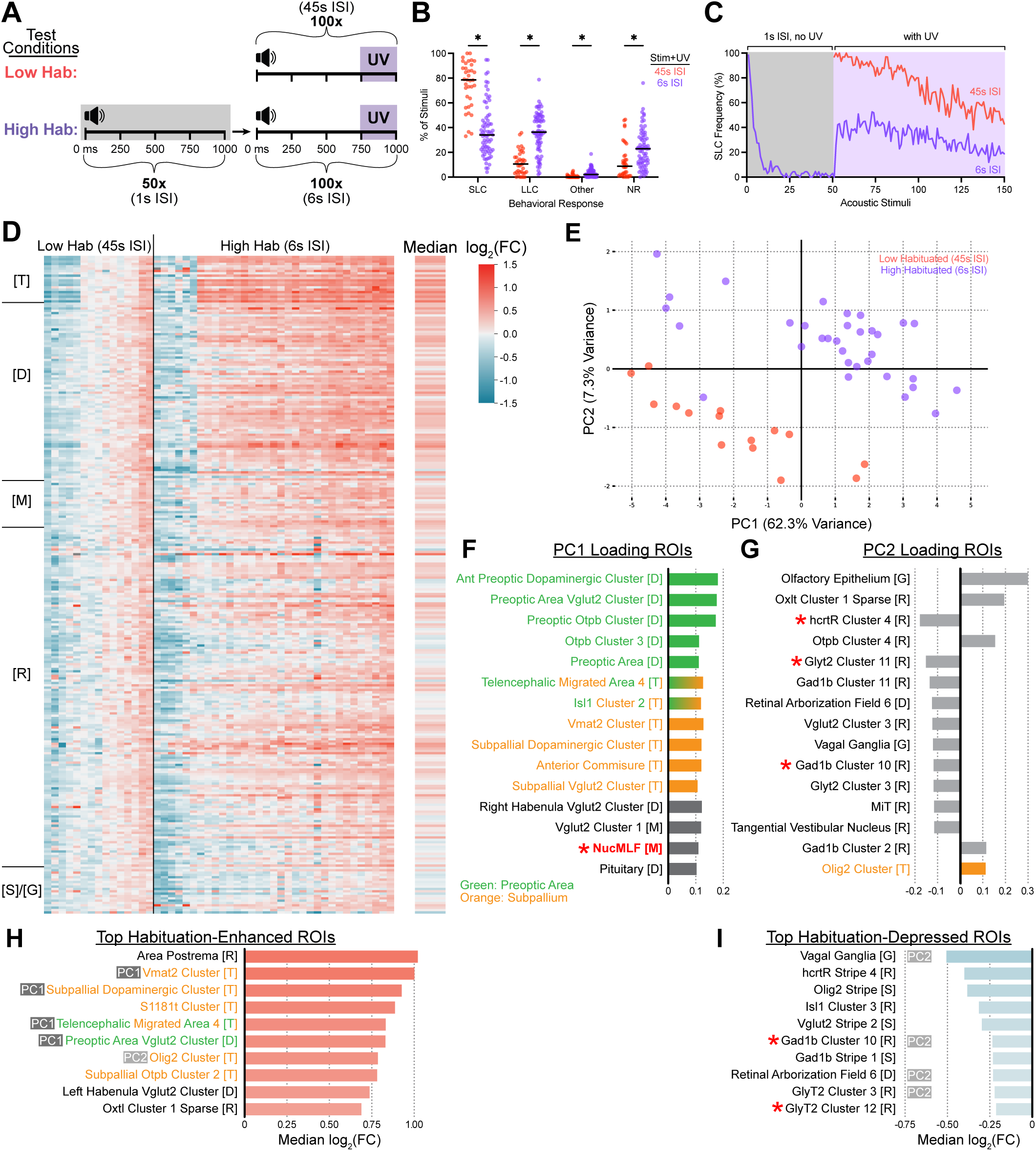
CaMPARI2 captures differential acoustically evoked neural activity between low- and high-habituating larvae. **(A)** Schematic of relative timings of acoustic stimuli and UV exposures. The low-habituating (Low Hab, top) condition consisted of 100 intense acoustic stimuli with a 45s ISI and accompanying 250-ms UV exposure beginning 750 ms after each acoustic stimulus onset as in Fig. 2A. The high-habituating (High Hab, bottom) condition consisted of 50 intense acoustic stimuli presented at 1 s ISI without accompanying UV exposure followed immediately by 100 intense acoustic stimuli presented at 6 s ISI with accompanying 250-ms UV exposures beginning 750 ms after each acoustic stimulus. **(B-C)** Frequencies of behavioral responses by 6 dpf wild type larvae to the 100 UV-coupled acoustic stimuli delivered in the Low Hab (red, n = 36 larvae) or High Hab (violet, n = 71 larvae). Larval behaviors were classified as Mauthner-dependent escapes (SLC), Mauthner-independent escapes (LLC), routine turn and swim bouts (Other), and no response (NR). Average frequencies of SLC responses across the entire experimental course are presented in (**C**). **(D)** Heatmap of log_2_(FC) in neural activity of larvae in low-habituating (45 s ISI, n = 15) and high-habituating (6 s ISI, n = 33) conditions relative to the low-habituating condition. Columns represent individual larvae and rows represent individual ROIs. The rightmost columns shows the median log_2_(FC) across all larvae in each post-stimulus condition. Colors corresponding to magnitude of log_2_(FC) are indicated by color bar. grouped by major brain regions along the y-axis: Telencephalon [T], Diencephalon [D], Mesencephalon [M], Rhombencephalon [R], and Spinal Cord and Ganglia [S]/[G]. ROIs with no detectable red or green fluorescence are shown in gray. Individual larvae and ROIs with >10% missing red or green fluorescence measurements were excluded from analysis. **(E)** Principal component analysis (PCA) of ROI-level RGRs. Each point represents one larva, with violet denoting to the high-habituating condition and red denoting the low-habituating condition. x- and y-axes show the first (PC1) and second (PC2) principal components, accounting for 62.3% and 7.3% of the total variance, respectively. **(F-G)** ROIs with the 15 largest loadings on PC1 (**F**) and PC2 (**G**). Bars lengths indicate loading value, and are color coded to reflect brain regions they are contained in (green: preoptic area, orange: subpallium). Asterisks indicate top acoustically-engaged regions identified in Fig 1**-3**. **(H-I)** ROIs displaying the ten largest median activity increases (**H**) and decreases (**I**) in the high-habituating condition relative to the low-habituating condition. Bars indicate median log_2_ (fold change) value. ROI name color indicates presence in the preoptic area (green) and/or subpallium (orange). ROIs loading to PC1 or PC2 are also indicated. Red asterisks indicate the ROI was among the top ten acoustically enhanced in Fig. 1**-3**.

We next directly compared the captured brain activities between the Low and High habituation conditions and observed distinct patterns of regional activity increases and decreases across high-habituating individuals relative to low-habituating individuals (**Fig. 4D**). Notably, broad patterns of relative regional increases and decreases appeared dissimilar to those observed in post-stimulus vs pre-stimulus conditions, with higher median activities in more rostral brain regions and less consistent activity increases in the rhombencephalon (**Fig. 4D**). Principal component analysis (PCA) of whole-brain patterns of ROI-level activity segregated high- and low-habituating larval groups into two distinct clusters separating along both PC1 and PC2, accounting for 62.3% and 7.3% of variance, respectively (**Fig. 4E**). This revealed robust and consistent multivariate activity differences between the high- and low-habituating conditions, establishing the two behavioral states as discrete (**Fig. 4E**). Examining PC1 and PC2 loadings identified the ROIs contributing most to discrete activity patterns between groups (**Fig. 4F-G**). PC1 was largely dominated by telencephalic and diencephalic ROIs, while PC2 was negatively loaded by many rhombencephalic ROIs, including several of the top ROIs from the previous low-habituating experiments (**Fig. 4G**, asterisks). Several ROIs loading most strongly into PC1 were also among those most differentially active in the positive direction (**Fig. 4H**). ROIs displaying the most significantly reduced activity in the high vs low habituating condition also overlapped with ROIs previously identified as having increased activity in stimulated larvae relative to no-acoustic or pre-acoustic UV capture larvae (**Fig. 4I**, red asterisks).

To further explore the neural basis of the non-associative learning process, we pharmacologically disrupted acoustically-evoked habituation learning by pretreating larvae with NMDA-type glutamate receptor (NMDAR) blocker MK-801 or vehicle alone for 30 minutes, then captured brain activities for both groups of larvae with a high-habituation stimulus paradigm (**Fig. 5A**). Similar to untreated larvae (**Fig. 4B**), vehicle-treated larvae were pre-shifted to a higher habituated state for the photoconversion phase with higher rates of non-response or LLC responses than SLC responses (**Fig 5B**, grey). In contrast, NMDAR-disrupted larvae largely responded to the same stimulus paradigm with Mauthner-dependent SLC responses and most individuals showed low frequencies of LLC responses or non-responses (**Fig. 5C**, green), reflecting their predicted habituation learning deficit (Wolman et al., 2011; Marsden and Granato, 2015; Nelson et al., 2023). Comparing brain-wide activity patterns between vehicle and MK-801 treated larvae revealed widespread relative reductions in activity across all broad structural regions of the brain (**Fig. 5D**), consistent with the global pharmacological disruption of excitatory glutamate neurotransmission. ROIs featuring the largest activity changes were largely rhombencephalic with reduced activity relative to vehicle-treated controls (**Fig. 5E**), and these included multiple ROIs that positively loaded to Habituation PC2 (**Fig. 5E**, purple). We specifically examined the ROIs driving Habituation PC1 to look for disrupted activation by MK-801, and all but the pituitary showed median activity reductions (**Fig. 5F**). Similarly, the top habituation-enhanced ROI’s all showed significant median fold changes in activity (**Fig 5G**), consistent with the behavioral reduction in habituation by MK-801. We also observed somewhat milder reductions in the median fold change of the top habituation-depressed ROIs, though these notably showed high variance between individuals **(Fig. 5H).** Together these data indicate that CaMPARI2 captures the neural activity changes associated with large-scale pharmacological circuit disruption.

**Figure 5.**
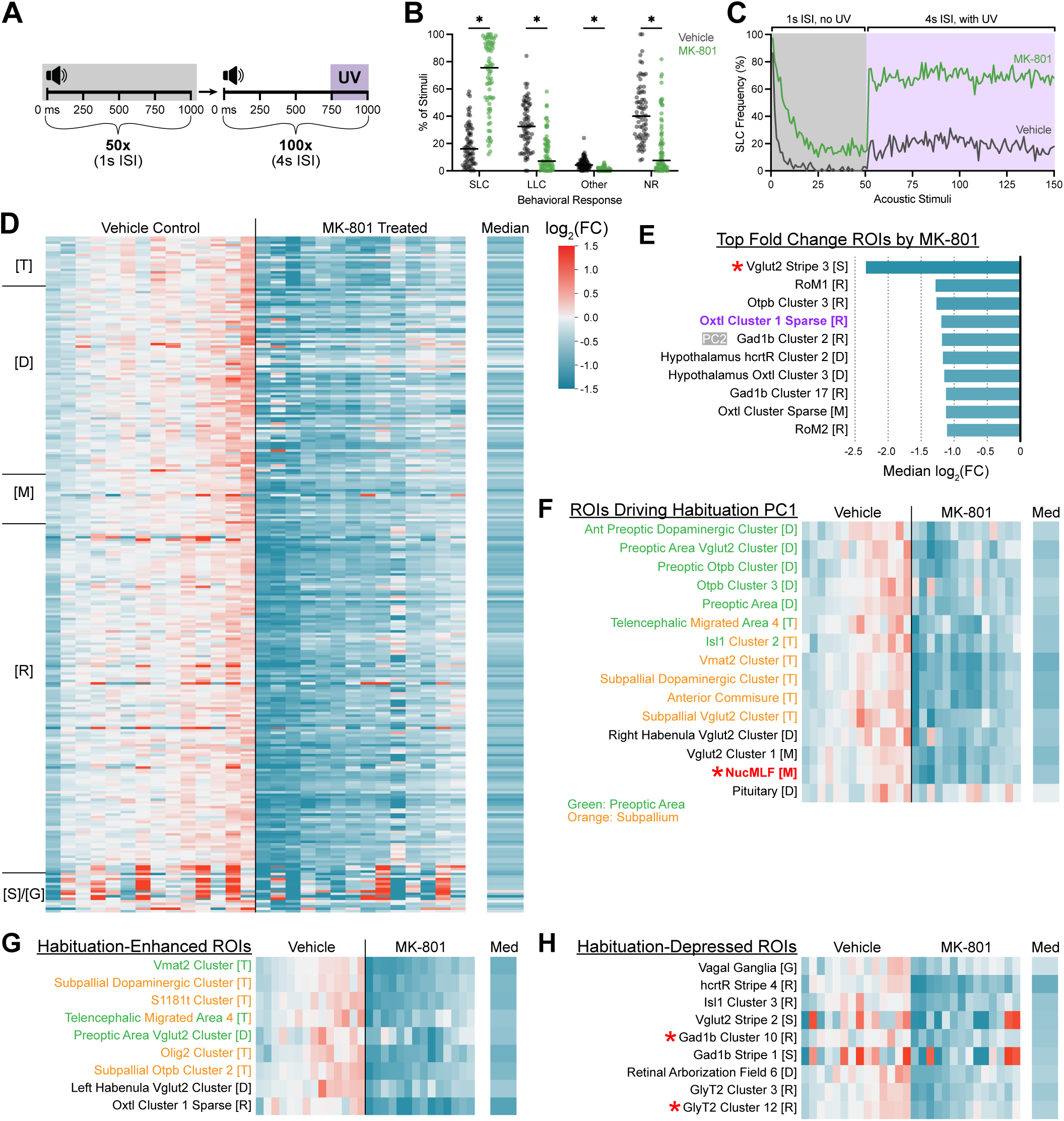
CaMPARI2 captures broad brain state differences when habituation is pharmacologically impaired by MK-801. **(A)** Schematic of relative timings of acoustic stimuli and UV exposures. Larvae pretreated for 30 minutes with vehicle or MK-801 were subjected to 50 intense acoustic stimuli presented at 1 s ISI without accompanying UV exposure followed immediately by 100 intense acoustic stimuli presented at 4 s ISI with accompanying 250-ms UV flashes beginning 750 ms after each acoustic stimulus. **(B-C)** Frequencies of behavioral responses by 6 dpf wild type larvae to the 100 UV-coupled acoustic stimuli delivered in the High Hab paradigm described in (**A**) following a 30-minute pretreatment with vehicle (gray, n = 72 larvae) or 500µM MK-801 (green, n-72). Larval behaviors were classified as Mauthner-dependent escapes (SLC), Mauthner-independent escapes (LLC), routine turn and swim bouts (Other), and no response (NR). Average frequencies of SLC responses across the entire experimental course are presented in (**C**). **(D)** Heatmap of log_2_(FC) in neural activity of larvae in vehicle-treated (n = 14 larvae) and MK-801-treated conditions (n = 14 larvae) relative to the median vehicle-treated condition for each ROI. Columns represent individual larvae and rows represent individual ROIs. The rightmost column shows the median log_2_(FC) across all larvae in the MK-801-treated condition. Colors corresponding to magnitude of log_2_ (fold change) are indicated by color bar. ROIs are grouped by major brain regions along the y-axis, Major brain regions are indicated on the left y-axis: Telencephalon (T), Diencephalon (D), Mesencephalon (M), Rhombencephalon (R), and Spinal Cord and Ganglia (SC & G). Log_2_(FC) values exceeding the displayed range are clipped and shown at the minimum of maximum color range. ROIs with no detectable red or green fluorescence are shown in gray. Individual larvae and ROIs with <10% missing red or green fluorescence measurements were excluded from analysis. **(E)** ROIs displaying the ten largest median activity decreases in the MK-801-treated condition relative to the vehicle-treated condition. Bars indicate median log_2_ (fold change) value. ROI in purple indicates it was a top Habituation-enhanced region in **Fig 4H**, and asterisks indicate overlap with top acoustically-engaged regions identified in Fig 1**-3**. **(F-H)** Heatmap subsets of log_2_(fold change) in neural activity of larvae in the MK-801-treated and vehicle-treated conditions from (**D**) including the identified ROIs contributing most strongly into habituation PC1 from Fig 4F (**F**), top 10 ROIs that were enhanced in high habituation conditions of normal fish in Fig. 4 **(G)**, and top 10 ROIs that were depressed in high habituation conditions of normal fish in Fig. 4 **(H)**. Columns represent individual larvae and rows represent individual ROIs. The rightmost column shows the median log_2_(FC) across all MK-801-treated larvae. ROI name color indicates presence in the preoptic area (green) and/or subpallium (orange). ROIs loading to PC1 or PC2 from are marked, and red asterisks indicate the ROI was among the top ten acoustically enhanced in Fig. 1**-3**.

## DISCUSSION

While visualizing neural activity is fundamental to understanding dynamic control of behavior, doing so with high temporal specificity in unrestrained animals proves challenging. Here we refine an approach to capturing temporally restricted, stimulus-locked neural activity in free-swimming larval zebrafish with CaMPARI2. By using the tool to investigate acoustically-evoked escape behavior, a temporally fixed paradigm initiated by spatially diverse neuroanatomical regions, we demonstrate its utility for elucidating differential activity patterns between larvae in various behavioral and learning states.

### Optimizing Key Variables for Temporally Specific CaMPARI2 Activity Capture

Selecting the appropriate timing and intervals of photoconverting UV light is crucial for isolating behaviorally relevant neural activity from background signal. Comparing individuals receiving 15x UV within-group in the absence or presence of acoustic stimuli revealed individual patterns with activity values diverging extensively from group medians (**Fig 1I**) suggesting substantial capture of irrelevant spontaneous neural activity and/or biological behavioral state differences (**Fig. 1I**). Consistent with this interpretation, acoustically relevant ROIs showed similarly variable activity patterns between replicates, with only modest and inconsistent activity increases in populations known to be critical for acoustically evoked escape (**Fig. 1G**). Identifying such activity patterns that are relevant can be enhanced by reducing noise from individual-to-individual variations where possible. We considered four possible sources of variation: 1) Acoustically-unrelated “baseline” neuronal activity, 2) Neural activity differences due to spontaneous motor behavior, 3) Neural activity differences related to acoustically-evoked motor behavior performance (ie SLC, LLC, NR), and 4) Neural activity differences in sensation & processing underlying behavior selection, learning, and memory. We typically sought to reduce the first two sources of variation for most paradigms, and the 3rd or 4th depending on the research question at hand. Despite identical acoustic stimuli between across conditions, modifying UV delivery parameters allowed us to capture and extract distinct regional activity patterns between experimental groups (**Fig. 1-3**).

#### Reducing the total cumulative UV

such as from 45 s to 22.5 s (**Fig. 1**), enhanced our ability to detect acoustically-evoked activity, and this should generally reduce the impact of acoustically-unrelated baseline and motor activities. We did not explore the lower limits of detection by CaMPARI2, though reducing from 25 s to 12.5 s total exposure still effectively detected our activity of interest albeit with (**Fig. 3**), though this was accompanied by reduced fold change values. While we predicted that acoustically-unrelated activity would be randomly sampled across UV exposure windows, this may still vary by internal state differences between individuals such as motivation, attention, hunger, or stress (Johnson et al., 2020; Krishnan et al., 2025; Légaré et al., 2025).

#### Shortening each UV window

such as from 3s to 1.5 s or 250 ms to 125 ms (**Fig 1-3**), reduced the likelihood of capturing spontaneous motor behavior in each window, since larval movements are performed in discrete discontinuous bouts (Marques et al., 2018). With acoustically-evoked escape behavior sufficiently small windows should exclude spontaneous behavior almost entirely, since performing escape bouts precludes other motor bouts (Burgess and Granato, 2007b). PC windows as small as 125 ms were sufficient to permit meaningful comparison between ROIs for acoustically-evoked escape behavior (**Fig. 3**), even without other compensation compared to 250 ms windows **(Fig. 2**), and we speculate that even shorter windows may also be effective and perhaps even distinguish between bout phases.

#### Increasing the number of UV-captured stimuli

such as from 15 to 100 (**Fig 1-2**), enhanced both the RGR of known acoustically-related ROIs captured and the magnitude of fold change of R/G over matched UV controls. Since CaMPARI2 integrates across captured epochs, increasing captured events enhances consistency when relevant behaviors aren’t performed 100% of the time (**Fig. 1F, 2B-C**), and increasing the number of captured stimuli can offset lower signal from short UV capture duration.

#### Increasing the ISI between UV-captured stimuli

as we did from 30 s to 45 s during piloting (**Fig 1A, E)**, increased the average SLC performance rate. In our “low habituation” condition overall SLC response rates significantly declined from stimulus 1 to 100 **(Fig 4C**). We anticipate that further increasing or altering the ISI would further reduce habituation to better distinguish different levels or periods of habituation learning (Wolman et al., 2011; Pantoja et al., 2016; Nelson et al., 2023). Since increasing numbers of UV-captured stimuli would also increase capture of “baseline” neural activity, it is important to balance this against potential other changes in behavioral states (Horstick et al., 2016).

#### Altering the relative timing of UV relative to stimuli of interest

such as shifting UV windows 250 ms prior to 750 ms following acoustic stimuli while holding other variables constant (**Fig 2, 3**), is critical to establish a window that closely matches the circuit function period of interest. Acoustically-evoked escape behaviors initiate in a restricted and stereotyped latency range (0-15 ms for SLC, 15-60 ms for LLC) and bout duration range, facilitating our UV window selection (Burgess and Granato, 2007b; Zúñiga Mouret et al., 2024). We found short windows of UV exposure optimally captured neuronal activity underlying acoustically-evoked motor behavior that had occurred 0.5-1 second *prior to the UV onset*, consistent with electrically-evoked activity in culture and slice preparations (Moeyaert et al., 2018; Perez-Alvarez et al., 2020). The 750-ms condition showed similar levels of PC, but with greater consistency between replicates across the whole brain (**Fig. 3C**) and an acoustically relevant ROI subset (**Fig. 3D**). ROIs displaying the largest stimulus-evoked activity increases remained fairly consistent between conditions, with those most active in the 625-ms condition overlapping most substantially with those previously identified by 250-ms UV exposure conditions (**Fig. 2E, 3E**). Taken together, our results point to a stimulus-to-UV delay time of 500-750 ms as optimal for acoustically-evoked activity capture.

#### Altering surrounding context of photoconversion periods

Altering the ISI between acoustic stimuli induces habituation learning, so comparing capture between different ISIs allows differentiating between learning state related activity. Similarly, introducing additional acoustic stimuli that were *not* accompanied by UV prior to the UV-captured stimuli (**Fig 4A, 5A**) shifted the behavioral profile larvae. Providing acoustic stimuli outside of the UV capture window for control individuals (**Fig 2, 3**) ensures that any larger scale behavioral state changes occurring for fish should be accounted for to focus on acute behavior selection and performance. Notably, excluding capture of interleaved or immediately prior stimuli is not possible with IEG-based approaches.

The CaMPARI2 system allowed us to directly compare stimulus-locked differential neural activity between larvae in naive and habituated behavioral states, comparing integrated neuronal activity states captured across 100 stimuli over 75 minutes vs 100 stimuli over 10 minutes (**Fig 4A**). In contrast, IEG- and pERK-based methods aggregate neural activity over the entirety of their extended detection windows, where detection is instead determined by the response and decay time course of the marker used, precluding comparisons between conditions in which the same number of stimuli are distributed over markedly different timescales. This temporal restriction difference allows the CaMPARI2-based approach to resolve both sustained, brain-wide changes associated with habituation and acute, stimulus-evoked differences in neuronal responses, despite the large differences in ISI and total experimental duration.

### Circuits Modulating Acoustically-Evoked Escape Behavior and Learning

Principal component analysis revealed that low- and high-habituated larvae form largely distinct clusters around PC1, which captured the majority of variance between the two stimulus delivery conditions (**Fig. 4E**). In conjunction with consistent patterns of activity increases and decreases localized to particular ROIs (**Fig. 4D**), this indicated that CaMPARI2 was able to resolve consistent, coordinated patterns of activity defining the two behavioral states.

To relate global separation observed in PCA to ROI-level habituation-driven activity, we examined the ROIs most strongly driving that separation, providing mechanistic insight into how whole-brain state differences emerge from region-level stimulus-evoked activity. Notably, several ROIs loading most strongly into the dominant principal component also ranked among the most strongly habituation-enhanced ROIs (**Fig. 4F, 4H**). The convergence of these analyses indicates that behavioral state separation is driven by coordinated upregulation of particular ROIs, pointing towards specific circuit elements selectively engaged during habituation. The overlap between ROIs loading most strongly into the second principal component and habituation-depressed ROIs suggests that these particular regions contribute to orthogonal, inter-individual variability in stimulus responses rather than driving primary behavior state separation (**Fig 4G, 4I**).

The ROIs most strongly segregating larvae by behavioral state and displaying the highest habituation-enhanced activity belonged to either the preoptic area or the subpallium (**Fig. 4F, 5F**), regions broadly connected to sensory adaptation and learning (Portavella et al., 2004; Lal et al., 2018; Nelson et al., 2023; Pandey et al., 2023; Palieri et al., 2024). The zebrafish preoptic area contributes to homeostatic navigation based on deviations from optimal water temperature, playing a vital role in behavioral modulation based on changing sensory context (Palieri et al., 2024). Though functionally heterogeneous, various regions of the subpallium have been implicated in fear and avoidance learning in teleost fish as well (Portavella et al., 2004; Lal et al., 2018; Pandey et al., 2023). In contrast to our results, habituation-deficient *pappaa* and *ap2s1* mutant larvae display consistent elevated subpallial and preoptic area activity over siblings (Nelson et al., 2023). Because these mutants show constant elevated pERK activity in these regions even without stimulation, it is possible this masked pERK-based assessment of temporally-specific recruitment of subpallial and preoptic area subpopulations we observed in wild type individuals through CaMPARI2, so more detailed cell- and time-resolved assessment of neurons promoting habituation within these regions is necessary to resolve the direct contributions of these neurons to habituation learning.

In contrast to the consistent and coordinated activity patterns observed in our habituation experiment, we noted that MK-801-treated larvae exhibited markedly increased replicate-to-replicate variability (**Fig. 5D**), indicating that CaMPARI2 captured the erosion of separation between habituated and non-habituated larvae following administration of MK-801. pERK-detection work has found that MK-801 dampens activity in the subpallium and habenula, which our findings replicated (**Fig. 5F, H**). Genetic disruption of neurotransmission in the zebrafish dorsal habenula impairs the modulation of responses to conditioned aversive stimuli, an associative learning process (Agetsuma et al., 2010). Additionally, dorsolateral-habenula-ablated juvenile zebrafish show a deficit in their ability to integrate past and novel experiences in a memory extinction and reversal learning context, positioning the region as vital for continuous behavioral modulation (Palumbo et al., 2020). Left dorsal habenula function is required for recovery of larval swimming behavior following a fear-inducing stimulus and subsequent response (Duboué et al., 2017). The lateral habenula has also been implicated in the transition between active and passive behavioral coping in larvae, demonstrating progressive encoding of sensory experience (Andalman et al., 2019).

Together, these observations demonstrate the relevance of preoptic, subpallial, and habenular populations in habituation learning and highlight the dependence of differential activity interpretation on the temporal specificity of capture. Future cell-level characterization of these regions will reveal their precise functionality in habituation learning.

### Opportunities and Limitations for Leveraging CaMPARI to Understand Behavioral Circuitry in Zebrafish

Our results demonstrate that brief windows of UV exposure optimally capture neuronal activity occurring 0.5-1 second prior to UV onset (Moeyaert et al., 2018; Perez-Alvarez et al., 2020). Both the UV-dependence and the delay inherent to this technique provide an opportunity for integration with closed-loop paradigms, in which behavioral responses influence the stimuli presented to an animal in real time (Jouary et al., 2016; Kawashima et al., 2016; Naumann et al., 2016; Vanwalleghem et al., 2018). Delivering photoconverting UV light contingent upon the execution of particular behaviors offers an avenue for investigating specific behavioral stages of complex and temporally variable scenarios, which would be difficult to capture with predetermined, temporally-fixed stimulus and UV delivery paradigms. Using this tool to provide mechanistic insight on other pharmacological or genetic variants shifting sensorimotor gating sensitivity or selection bias between escape responses, where capturing temporally precise subsets of neural activity is essential (Burgess and Granato, 2007b; Wolman et al., 2015; Jain et al., 2018; Marsden et al., 2018). Combining zebrafish whole brain CaMPARI2 studies with genetic variants linked to human neuropsychiatric conditions would facilitate mechanistic neurobiological understanding, though assessing neurodevelopmental structural differences will be critical to appropriately interface with standardized brain atlases (Gupta et al., 2018; Thyme et al., 2019; Marquez-Legorreta et al., 2022; Mendes et al., 2023; Capps et al., 2025). While the temporal specificity and stereotyped nature of acoustically-evoked behavior make CaMPARI2 an effective tool to dissect the underlying neural activity, this pipeline could be implemented to investigate behaviors engaging other sensory modalities where restraint can compromise behavioral features, such as hunting or social behaviors (Tunbak et al., 2020; Zhao et al., 2025).

At the same time, the dependence of CaMPARI2 activity capture on UV light introduces important considerations. Zebrafish larvae exhibit high spectral sensitivity to UV light and display intensity-dependent avoidance behavior, raising the possibility that UV exposure itself could disrupt visually guided behaviors, influence multisensory integration, or act as an unintended associative learning cue (Nava et al., 2011; Guggiana-Nilo and Engert, 2016). Indeed, during our piloting experiments we noted altered responsiveness and habituation rates when UV flashes were incorporated into the stimulus paradigm at different points relative to acoustic stimuli. We also observed that sufficiently short UV pulses fail to evoke light flash induced responses during piloting, suggesting UV photoconversion doesn’t necessarily preclude visually-guided behavior analysis through CaMPARI2, consistent with its recent use capturing mouse visually-guided behaviors (Burgess and Granato, 2007a; Das et al., 2023a). These observations underscore the necessity of incorporating UV flashes into reference conditions and behavioral assay optimization.

Combining CaMPARI2 with complementary experimental approaches offers the opportunity to link whole-brain activity patterns to subcellular activity dynamics and molecular determinants of circuit function. Incorporation of pre- and post-synaptically targeted CaMPARI2 variants like SynTagMA would permit activity capture at the synaptic level, enabling finer dissection of particularly distributed circuits spanning large brain areas (Perez-Alvarez et al., 2020). Recent work in both mice and larval zebrafish has also combined CaMPARI-based activity capture with fluorescence-activated cell sorting and subsequent transcriptomic profiling, allowing populations recruited during particular behavioral states to be characterized and linking circuit dynamics to gene expression programs (O’Toole et al., 2023; Matsuda et al., 2025). Both CaMPARI2 whole-brain activity maps and transcriptomic datasets derived from CaMPARI2-labeled neurons lend themselves to a variety of computational analyses. Applied to whole-brain activity, multivariate, clustering, and network-based approaches can uncover coordinated patterns of circuit engagement and behavioral-state-dependent differences in whole-brain activity, as demonstrated by distinct clustering of habituation states in principal component space (Constantin et al., 2020; Marquez-Legorreta et al., 2022; Légaré et al., 2025). In contrast, the absence of comparable clustering of pharmacologically-induced behavior states highlights how such analyses can reveal more heterogenous, globally dysregulated activity patterns, consistent with the variable and widespread effects of NMDA receptor antagonism (Moghaddam et al., 1997; Homayoun and Moghaddam, 2007). Additionally, different experimental perturbations require tailored computational frameworks to faithfully assess their impact on neural activity (Cunningham and Yu, 2014). Altogether, expanding the zebrafish toolkit for whole-brain circuit activity analyses using CaMPARI2 effectively complements existing methods and expands the accessibility of large scale behavioral circuit dissection beyond highly specialized real-time volumetric imaging equipment.

## MATERIALS AND METHODS

### Zebrafish Maintenance and Husbandry

Zebrafish maintenance, husbandry, and raising of larvae was performed in accordance with standard published guidelines. All experimental protocols were approved by the Haverford College IACUC. Zebrafish were maintained on a 14h light/10h dark cycle. Embryos and larvae were raised in 1x E3 (5mM NaCl, 0.17mM KCl, 0.33mM CaCl_2_, 0.33mM MgSO_4_) at 29°C. Larvae used for whole-brain confocal imaging were raised in 200 µM phenylthiourea (PTU, Sigma, P-7629) in 1x E3 beginning at 24 hpf to inhibit melanophore development. Embryos were treated with 20 µg/mL pronase (Sigma, 10165921001) between 20 and 28 hpf to facilitate synchronous dechorionation. Pronase was removed and replaced with fresh PTU/E3 after 24 h of exposure. Prior to imaging, larvae were screened for central nervous system CaMPARI2 fluorescence and developmental abnormalities. Larvae between 5 and 7 dpf were used for all imaging and behavioral experiments. As there are currently no consistent known genetic markers for sex in zebrafish and sexual differentiation does not occur until 25-60 dpf, all larvae were tested without regards to sex. Larvae were humanely euthanized via rapid chilling or tricaine overanesthesia following experiments in accordance with AVMA guidelines. All CaMPARI2 imaging experiments presented in this manuscript used the *Tg(elavl3:CaMPARI2)jf92* transgenic line, referred to as *HuC:CaMPARI2* (ZDB-ALT-180410-5), though some original pilot experiments were performed using the earlier *Tg[elavl3:CaMPARI(W391F+V398L)]jf9* line (ZDB-ALT-150403-1). Transgenes were maintained in the Tüpfel long fin (TLF) background (ZDB-GENO-990623-2) carrying a *mitfa* mutation (ZDB-ALT-060913-2) to reduce skin pigmentation for enhanced imaging quality. For all behavioral assays, the TLF strain was used with normal pigmentation to facilitate automated behavioral tracking.

### CaMPARI2 Photoconversion and Acoustic Behavior Assays

For photoconversion, four 5-7 dpf larvae were placed into one 9 mm diameter circular well of a 36-well custom plexiglass arena attached to a vibrational exciter (4810, Brüel & Kjær) delivering non-directional vibroacoustic stimuli. Acoustic stimuli were administered using DAQTimer software (Burgess and Granato, 2007a) at a frequency of 1000 Hz, intensity of 23.5 dB, and duration of 2 ms, as previously described (Zúñiga Mouret et al., 2024), and at the intervals specified in the assay diagrams presented (**Fig. 1A, 1E, 2A, 3A, 4A, 5A**). UV light was delivered via a lightguide-coupled LED light source (UHP-T-405-DI, Prizmatix) with a spectral maximum of 405 nm, and was also controlled via DAQTimer. For photoconversion experiments, the UV light source was placed directly above the well containing the larvae, approximately 2 cm from the surface of the water to provide consistent illumination.

Since the photoconversion UV light source occluded larvae from overhead video recording, in our representative behavioral analyses, the UV light was presented obliquely from above approximately 20cm from the surface such that UV light would fully illuminate all wells containing fish. A high-speed camera (TS4, Fastec) fitted with a Sigma 50mm f/2.8 EX DG Macro Autofocus lens (B&H Photo) and 720 nm wavelength filter (Opteka) filter was used to record larval behavior at 1000 fps. An infrared array (CM-IR200-850, CMVision) provided illumination for the camera below the white plexiglass diffuser base of the testing arena, as previously described (Zúñiga Mouret et al., 2024). For the representative behavioral analyses, each 9mm well either contained four wild type TLF larvae (**Fig. 1E**) or individual wild type TLF larvae (**Fig. 2-5**) raised without PTU, unlike larvae destined for photoconversion, to preserve normal body pigment and permit automated behavioral tracking via the Flote software package (Burgess and Granato, 2007a). All behavioral experiments were carried out at 6 or 7 dpf, and compared between larval siblings.

### Pharmacological Treatment of Larvae

NMDAR glutamate neurotransmission was acutely disrupted as previously described (Wolman et al., 2011; Nelson et al., 2023). A 100 mM stock of MK-801 was prepared by dissolving powdered (+)-MK-801 hydrogen maleate (Sigma Aldrich, M107) in 100% DMSO. Prior to photoconversion, five 5-7 dpf larvae were placed into one 21.9 mm x 17.5 mm (diameter x height) circular well of a standard 12-well plate containing 2 ml of 1x E3. Thirty minutes prior to UV and stimulus delivery, 10 μL of either 100% DMSO or 100 mM MK-801 was added to the wells for final concentrations of 0.5% DMSO and 500 μΜ MK-801.

### Whole-Brain Imaging of Larvae

Following photoconversion, larvae were anesthetized with buffered 0.01-0.015% tricaine (Syndel) and mounted dorsally in 80 μL 1.5% low-melting point agarose (Sigma, A9414) on 35 mm coverslip bottom dishes (Mattek) and overlaid with diluted tricaine/E3. Larvae were mounted with the forebrain and optic tectum flush against the slipcover with minimal lateral tilt. Whole-brain, multichannel z-stacks were collected on a Leica Stellaris 5 line-scanning confocal microscope using a 20x water immersion objective. Unconverted CaMPARI2 fluorescence was collected in the green channel with 488 nm excitation at 40.08% intensity and a detection range of 495-554 nm with a gain of 33.0% (Moeyaert et al., 2018). Photoconverted CaMPARI2 fluorescence was collected in the red channel with 561 nm excitation at 90.08% intensity and a detection range 566 nm and 700 nm with a gain of 70.0% (Moeyaert et al., 2018). Two overlapping 512x512 z-stacks were collected for each brain (1 µm Z-step size, bidirectional scanning, collecting green then red channels sequentially for each 512x512 stack) and automatically stitched together through the Leica LAS X software. Z-stack endpoints were set just below and above the most dorsal and ventral portions of the brain, respectively, to capture the entire brain at a voxel resolution of 1.14 µm x 1.14 µm x 1.0 µm. For representative brain image presentation (**Fig. 1 B-D**), image stacks were captured at identical settings, and the photoconverted channel was isolated and switched to the Fire LUT before Z-projection in Fiji/ImageJ (Schindelin et al., 2012).

### ROI-Based Fluorescence Quantification

Pre-processing, registration, and fluorescence quantification of whole-brain images was based on the Multivariate Analysis of Variegated Expression in Neurons protocol (Shoenhard and Granato, 2023). Merged multichannel z-stacks in LIF (Leica image file) format were opened in Fiji/ImageJ (Schindelin et al., 2012) and reoriented such that the forebrain pointed upwards. The red and green channels were separated and stack-sorted such that images were ordered from the ventral to dorsal direction, and saved individually as NRRD files with suffixes denoting each channel. Files were then registered to the Z-brain atlas ZBB_jf9_huC-CaMPARI reference brain with the Computational Morphometry Toolkit (CMTK) Fiji plug-in for Mac OS (Randlett et al., 2015; Shoenhard and Granato, 2023).

Registered NRRD files were downsampled, smoothed, and converted to TIFF files using the PrepareStacksForMAPMapping.ijm Fiji macro (Randlett et al., 2015). Red and green fluorescence was quantified in each of 293 neuroanatomical regions (as defined by the Z-brain atlas) with the QuantifySignalMultipleBrains.m MATLAB script, which uses the MaskDatabaseDownsampled.mat and AnatomyLabelDatabase.hdf5 reference files (Randlett et al., 2015).

Regional fluorescence quantification returns a CSV file of red and green fluorescence in each ROI for each larva. All subsequent data processing was performed in Python (v3.14) with pandas and NumPy. Red fluorescence was normalized to green fluorescence by calculating a red-to-green ratio (RGR) for each ROI and larva. For each ROI, the median RGR was calculated across control replicates to capture activity unrelated to the behavior of interest as a reference. RGR values from experimental and control replicates were divided by the corresponding median value of the control population to calculate fold activity changes. For all downstream analyses and visualizations, ROIs and larvae with >10% of fluorescence values missing were excluded. We excluded the following ROIs from all of our analyses presented here because their superficial positions prevented consistent or reliable fluorescence readings across warped brains: facial glossopharyngeal ganglion, posterior and anterior lateral line ganglia, and lateral line neuromasts D1, D2, N, O1, OC1, SO1, SO2, and SO3. We also excluded the Mauthner neurons ROI from presented data, as these neurons do not appear to reliably express CaMPARI2 in all fish expressing the *jf92* transgenic line used.

### Data Visualization and Statistical Analyses

All data visualization was conducted in Python (v3.14) using Matplotlib, seaborn, and scikit-learn. To generate heat maps, fold changes were log_2_ transformed to render activity increases and decreases symmetrical and enable comparison between ROIs. To aid in visualization, Log_2_(fold change) values were thresholded at -1.5 and 1.5 in heat maps.

PCA was performed on mean-centered ROI-level RGR values with individual larvae treated as observations and ROIs as features. Values missing after data filtering were imputed using the mean value for each ROI. The first two principal components were computed and visualized to identify ROIs contributing most strongly to each component.

## AUTHOR CONTRIBUTIONS

KRR: Conceptualization, Data curation, Formal analysis, Investigation, Methodology, Project administration, Software, Validation, Visualization, Writing – original draft, Writing – Review & Editing

AB: Formal analysis, Investigation, Writing – Review & Editing;

RAO: Conceptualization, Formal analysis, Investigation, Methodology, Writing – Review & Editing;

KSV: Investigation, Writing – Review & Editing;

EES: Investigation, Writing – Review & Editing;

EM: Formal analysis, Investigation, Writing – Review & Editing;

AC: Formal analysis, Investigation, Visualization, Writing – Review & Editing;

PBD: Formal analysis, Investigation, Visualization, Writing – Review & Editing;

CC: Formal analysis, Investigation, Visualization; Writing – Review & Editing;

GCP: Conceptualization, Formal Analysis, Methodology, Software, Writing – Review & Editing;

RAJ: Conceptualization, Formal analysis, Funding acquisition, Investigation, Methodology, Project administration, Resources, Supervision, Validation, Visualization, Writing – Review & Editing.

## ACKNOWLEDGMENTS

We would like to thank Dr. Zak Kerrigan for instrument support, Laura McRae for oversight of the Haverford College Zebrafish Facility, and Nicole Cunningham for reagent support, Dr. Hannah Shoenhard and Dr. Jessica Nelson for sharing code and helpful discussion, and Dr. Kurt Marsden for providing the *HuC:CaMPARI2* transgenic line.

## DATA AVAILABILITY STATEMENT

Datasets and code are available on request from the corresponding author.

